# Fluorescence-Activated Droplet Sorting of Polyethylene Terephthalate-degrading Enzymes

**DOI:** 10.1101/2021.05.12.443719

**Authors:** Yuxin Qiao, Ran Hu, Dongwei Chen, Li Wang, Ye Fu, Chunli Li, Zhiyang Dong, Yunxuan Weng, Wenbin Du

## Abstract

Enzymes that can decompose synthetic plastics such as polyethylene terephthalate (PET) are urgently needed. However, a bottleneck remains due to a lack of techniques for detecting and sorting environmental microorganisms with vast diversity and abundance. Here, we developed a fluorescence-activated droplet sorting (FADS) pipeline for high-throughput screening of PET-degrading microorganisms or enzymes (PETases). The pipeline comprises three steps: generation and incubation of droplets encapsulating single cells, picoinjection of fluorescein dibenzoate (FDBz) as the fluorogenic probe, and screening of droplets to obtain PET-degrading cells. We characterized critical factors associated with this method, including specificity and sensitivity for discriminating PETase from other enzymes. We then optimized its performance and compatibility with environmental samples. The system was used to screen a wastewater sample from a PET textile mill. We successfully obtained PET-degrading species from nine different genera. Moreover, two putative PETases from isolates *Kineococcus endophyticus* Un-5 and *Staphylococcus epidermidis* Un-C2-8 were genetically derived, heterologously expressed, and preliminarily validated for PET-degrading activities. We speculate that the FADS pipeline can be widely adopted to discover new PET-degrading microorganisms and enzymes in various environments and may be utilized in the directed evolution of PETases using synthetic biology.

## Introduction

Synthetic plastics have become an indispensable part of modern life. The problem of plastic pollution is intensifying, especially after the COVID-19 (Coronavirus disease 2019) pandemic broke out^1^. Due to its excellent chemical durability and thermal properties, polyethylene terephthalate (PET) has become the primary thermoplastic resin for making bottles and fibers^2^. The control, recycling, and treatment of PET wastes is a great challenge. As a result, significant interests have arisen in unveiling the biodegradation of PET by microorganisms to tackle the problems caused by PET wastes^3^. Only a few microbial PET-degrading enzymes (PETases) have been reported, including cutinase-like enzymes from Thermobifida^4-5^ and lipases or esterases from Candida antarctica^6^ and Yarrowia lipolytica^7^. A PETase derived from Ideonella sakaiensis has been found that degrades PET into mono(2-hydroxyethyl) terephthalate (MHET) and terephthalic acid (TPA)^3, 8^. DuraPETase^9^, EXO-PETase^10^, and leaf-branch compost cutinase (LCC)^11^ were recently developed by structure-based protein engineering to improve catalytic activity and thermostability. Despite all these breakthroughs, we expect that much more microbial PETases have not yet been discovered based on the great genetic diversity and abundance of microorganisms in nature. However, discovering new PETases from environment is time-consuming and labor intensive^12^. It involves enrichment, screening, cultivation, enzyme expression, and activity validation. The screening of novel degrading microbes and enzymes remains a slow and complex process due to the inefficiency of current sorting techniques^5, 13-14^. Traditional measurements for plastic biodegradation include plastic weight loss, changes in the mechanical properties or the chemical structure, and carbon dioxide emission, but these can take up to several months. High-performance liquid chromatography (HPLC) or absorbance assays have been developed for PET film hydrolysis^15^ but have not yet been scaled up for high-throughput analysis of environmental microbial communities or large-scale mutant libraries. It is still difficult to efficiently evaluate the activity of PETases due to the lack of rapid, specific, sensitive, and quantitative detecting and sorting methods.

To overcome the bottleneck for the screening of functional microbes or enzymes, various high throughput screening techniques have been developed^16-17^. Among them, microfluidic fluorescence-activated droplet sorting (FADS) was introduced to dramatically simplify the operations, improve sorting efficiency and flexibility, and reduce the cost of large library screening^18-19^. FADS has been successfully applied to screen lipase/esterase^20^, DNA/XNA polymerase^21^, cellulase^22^, and NAD(P)-dependent oxidoreductases^23^ by incorporating highly sensitive and specific fluorescent enzymatic assays^24^. In this work, we developed a FADS-based approach for expediting the search of PETases and then validated its performance by benchmark PETases. We used it to obtain PET-degrading microbes from the wastewater of a PET textile mill, and evaluated putative new PETases for PET-degrading activities. We envision that the FADS pipeline developed in this study can be widely applied to the discovery of PET-degrading microorganisms and enzymes.

## Results

### FADS Pipeline for PETases

We established a complete FADS pipeline for sorting of PET-degrading microbes (PETases) consisting of four consecutive steps (**Fig. 1A**): **(i)** The microbial suspension is separated as single cells in picoliter droplets by droplet maker device. PET-degrading species were provided with a suitable long-term incubation without interference from a competing species. **(ii)** The droplets were re-injected into the picoinjection device to introduce picoliter fluorescein dibenzoate (FDBz) into each droplet and incubated at room temperature for a short time to allow the facilitate of FDBz. **(iii)** The droplets were re-injected into the sorting device for high-throughput sorting to obtain microbial species capable of decomposing FDBz (the fluorogenic probe for PETases). **(iv)** The positive droplets were then demulsified, and the positive species were isolated and identified on BHET agar plates. Afterward, The PET degradation performances of positive isolates were evaluated by fermentation with PET fibers and observation of surface erosion of PET films.

**Fig. 1.**
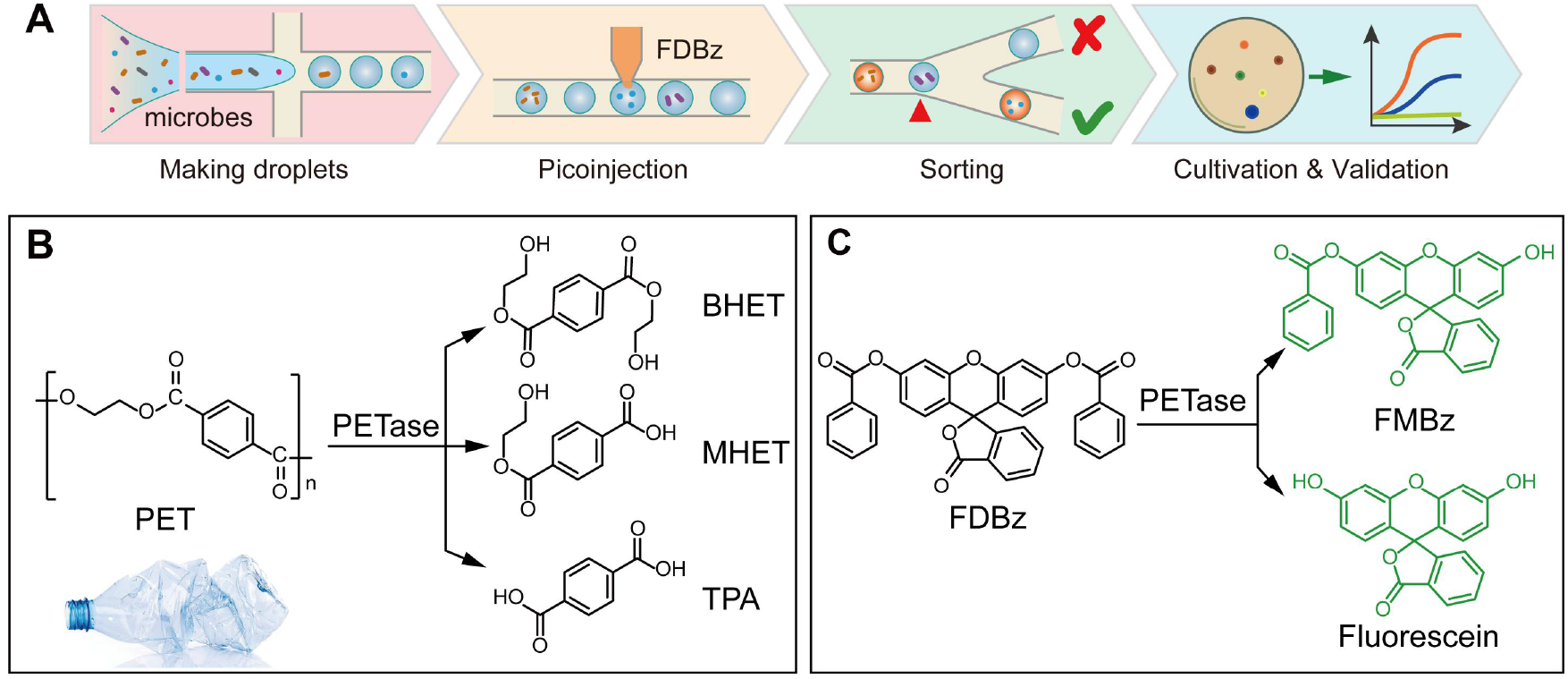
The workflow for sorting PET-degrading microbes by droplet microfluidics using fluorescein dibenzoate (FDBz) as the fluorogenic substrate. (A) Sorting procedures for isolation, cultivation and validation of PET-degrading microbes. (B) Biodegradation of PET by PETases releases MHET, BHET, and TPA. (C) Hydrolysis of FDBz releases FMBz and fluorescein.

### Fluorogenic Assays for PETases in Picoliter Droplets

To develop a rapid, sensitive, and specific fluorogenic assay for droplet-based single-cell sorting of PETases, the following criteria have to be met: (a) structural similarity between the fluorogenic probe and PET polymer structure; (b) good sensitivity and specificity for fluorescence discrimination of PETases from other enzymes; and (c) low level of self-hydrolysis, leakage and cross-talk of the fluorogenic probe and its hydrolyzed products between droplets. Most microbial-oriented PETases catalyze the hydrolysis of ester bonds of PET next to the benzene rings (**Fig. 1B**). In criteria (b), we selected FDBz as the candidate fluorogenic probe and evaluated its specificity and sensitivity for fluorescence detection of PETases in picoliter droplets. As previously reported, FDBz was not a highly reactive substrate of common lipases and cannot not be used to stain cancer cells or plant pollen cells^25^. However, FDBz has PET-like ester bonds linked with a benzene group and can likely be hydrolyzed by PETases to generate fluorescein monobenzoate (FMBz) and fluorescein (**Fig. 1C**), which can be detected by fluorescence. We used benchmark PETase and lipase to test the specificity of FDBz in selective reaction with PET-degrading enzyme. We performed colorimetric assays, fluorometric analysis, and the droplet-based catalytic reaction of benchmark PETases using FDBz as the substrate. The colorimetric assays were performed in Eppendorf tubes by mixing 250 μM FDBz with various enzyme solutions separately including cutinase (10 μM, a positive PETase), lipase (10 μM, negative control), and Tris-HCl buffer (blank control). After two hours of incubation, only the tube with cutinase turned yellow and fluorescent, while the tubes with lipase and the control remained unchanged (see **Fig. S3**). Next, quantitative fluorometric analyses were performed using the same reaction setting on a plate reader at an excitation wavelength of 488 nm and an emission wavelength of 523 nm. During a 2-h incubation at 37°C, the fluorescence intensity of microwells with cutinase (green line) increased nearly 346.5-fold versus 19.9-fold increase with lipase (orange line); the blank control (grey line) remained flat during 2-h incubation indicating high specificity and substrate selectivity of cutinase-FDBz hydrolysis (**Fig. 2A**). We further studied the effect of cutinase concentration on FDBz hydrolysis. The results shows that the fitted dose-response curve agrees well with classical Michaelis-Menten kinetics (R^2^=1) (**Fig. 2B**).

**Fig. 2.**
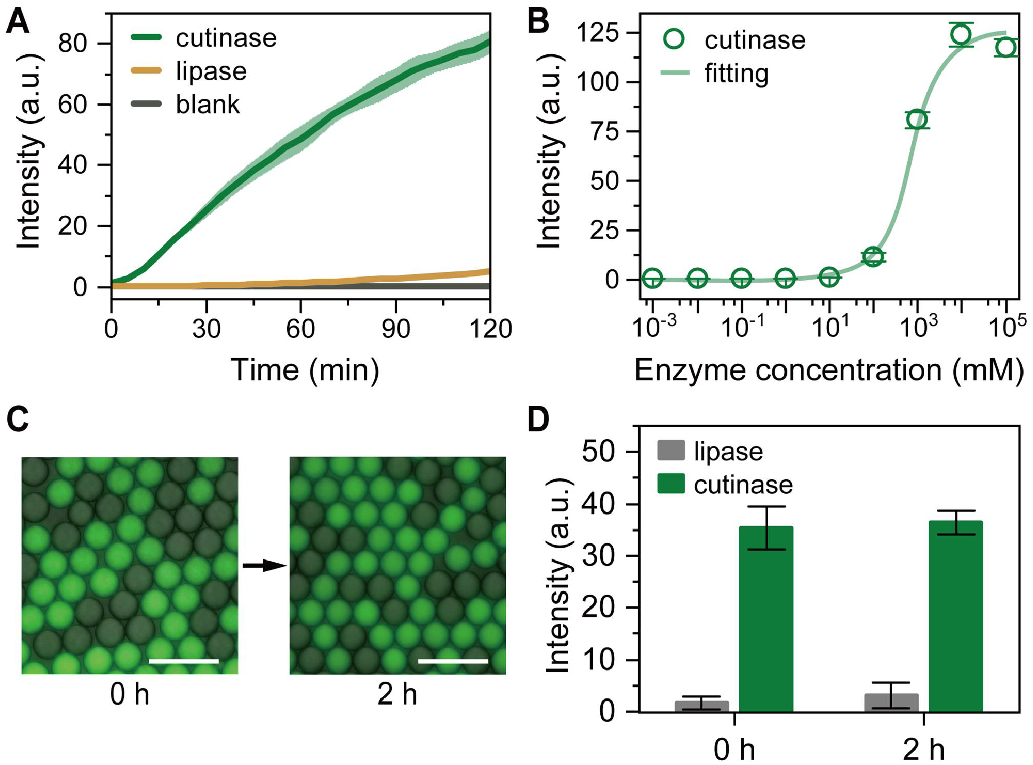
The FDBz-based fluorogenic assays for PETases. (A) The fluorescence reaction in the microplate. (B) Specificity and selectivity of cutinase-FDBz hydrolysis comparing with lipase and blank control. (C) FDBz-based fluorogenic assays in 4-pL droplets were indicating high sensitivity, low leakage, and low self-hydrolysis. (D) Fluorescence intensity difference between cutinase-FDBz and lipase-FDBz droplets before and after 2-h incubation. (Scale bar: 50 µm)

To evaluate whether FDBz is suitable for droplet-based PETase screening, we performed assays by mixing FDBz with either cutinase or lipase to make positive and negative droplets at a volume of 4 pL separately with yield final concentrations of 125-μM FDBz and 5-μM enzyme. First, we used the FADS optical system to read positive or negative droplets incubated separately for 40 minutes at the same settings. The histogram showed that fluorescence intensities of positive droplets were in the range of 2.0×10^6^ to 8.5×10^6^, and the fluorescence intensities of negative droplets were in a much lower range of 1.0×10^4^ to 5.0×10^4^ (**Fig. S4**). The difference of fluorescence intensities allows absolute threshold setting to discriminate PETases from common lipases and a sorting throughput of 1000 droplets per second with sorting efficiency of close to 100% using our FADS system. Afterward, positive and negative droplets were mixed, and monolayer arrays of droplets were prepared for time-lapse fluorescence imaging to investigate the leakage and self-hydrolysis (**Fig. 2C**). The ratio of fluorescence intensities of positive droplets against negative droplets decreased from 20.88 to 11.49 after 2-h co-incubation (**Fig. 2D**), but this is still sufficient for absolute discrimination of droplets. Overall, similar results were found in both bulk and droplet-based assays, proving that FDBz is qualified as a specific fluorogenic substrate of PETases in our FADS pipeline.

### FADS of Environmental PET-degrading Microbes

We applied the FADS pipeline to screen PET-degrading microbes from the wastewater of a PET textile mill located in Shaoxing city (Zhejiang, China). Bacterial suspensions of the original samples and its enrichment culture were screened to discover new PET-degrading microbes and PETases (**Fig. 3A, 3B**). The FADS was performed at a throughput of 1000 drops per second for several hours. As we expected, after enrichment cultivation, the positive rates with droplet fluorescence intensity above the sorting threshold increased to 2.98% versus 0.27% for the original samples using the same conditions (**Fig. 3A, B**). The sorted positive droplets were then demulsified and subjected for cultivation on agar plates. In total, we obtained 17 potential PET-degrading isolates belonging to eight genera, including nine isolates from the original sample and six isolates from the PET-YSV enrichment (**Table S1**).

**Fig. 3.**
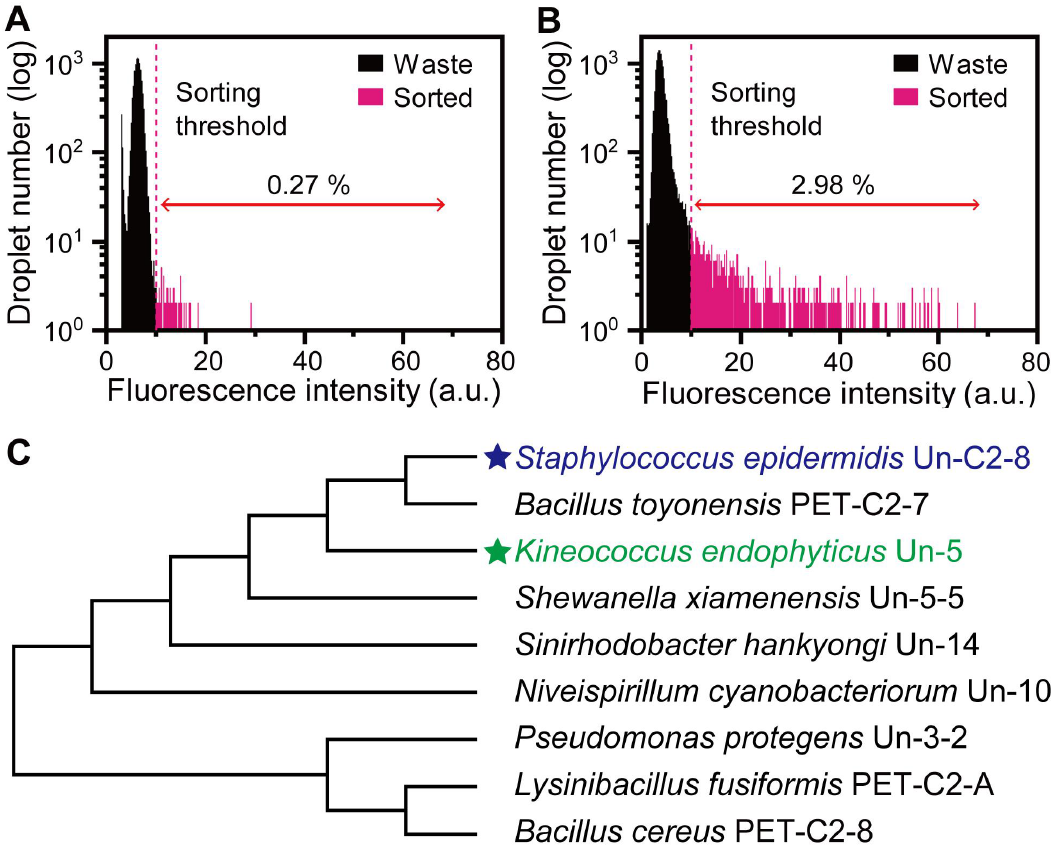
FADS of PET-degrading microbes from wastewater of a PET textile mill. Histograms showing droplet fluorescence of the original (A) and PET-YSV enriched (B) cell suspensions. The pink dashed lines indicate the sorting threshold. (C) The phylogenetic tree of PET-degrading strains obtained by FADS from a PET textile mill wastewater.

Interestingly, the isolated species from the original sample were different from those obtained from the PET-YSV enrichment and indicate that the enrichment cultivation might lead to the rapid growth of fast-growing opportunistic strains and suppressed those slow-growing PET-degrading species in the original sample.

We used the hydrolysis of BHET on agar plates to preliminarily evaluate the degrading activity of obtained microbial strains. BHET hydrolysis activity was determined by forming clear zones around the punched holes on both BHET agar plates. Among the nine strains that tested positive for clear-zone formation (**Fig. 3C, Table S1**), *Kineococcus endophyticus* Un-5 and *Staphylococcus epidermidis* Un-C2-8 exhibited the highest degrading activity and were selected for further degradation evaluation.

To further confirm the degradation activity, PET films and fibers were chosen as the degrading materials (**Fig. S5**). Isolates Un-C2-8 and Un-5 were inoculated in 20 mL YSV medium containing 40 mg PET films, and cultivated at 37°C. After two weeks, the hydrolyzed products, including BHET, MHET, and TPA, were quantified by HPLC. The HPLC profiles revealed that TPA and MHET in Un-C2-8 and Un-5 cultures were much higher than those of *E. coli* and blank control. The total amounts of released products (including TPA and MHET) were as high as 16.68 μg and 1.55 mg for Un-C2-8 and Un-5, respectively (**Figure4 A, B**). Meanwhile, similar degradation activities were also observed for both strains when cultivated with 60 mg PET fibers serve as the carbon source in 20 mL YSV medium for two weeks (**Fig. S6**). The yields of TPA were 37.63±1.45 μg and 50.98±3.70 μg for Un-C2-8 and Un-5, and MHET were 14.39±0.96 μg, and 16.20±7.54 μg for Un-C2-8 and Un-5, respectively.

To further confirm the biodegradation of PET by the microbial isolates using SEM imaging, the PET films were recovered from the flasks cultivated with Un-C2-8, Un-5, and the controls. **Fig. 4C shows** distinctly mottled surface erosion of PET films by strains Un-C2-8 and Un-5 versus the smooth surface of PET film incubated with *E. coli* or blank YSV medium. Un-C2-8 displayed an immersive erosion pattern, which is more likely to secret enzymes to degrade PET film. In some cases, the attached Un-C2-8 cells caused surface erosion and broke the PET film into porous fragments (**Fig. S7**). In contrast, Un-5 showed regional surface erosion that might form bacterial biofilms attached with the surface and thus create erosion hot spots (**Fig. 4C**).

**Fig. 4.**
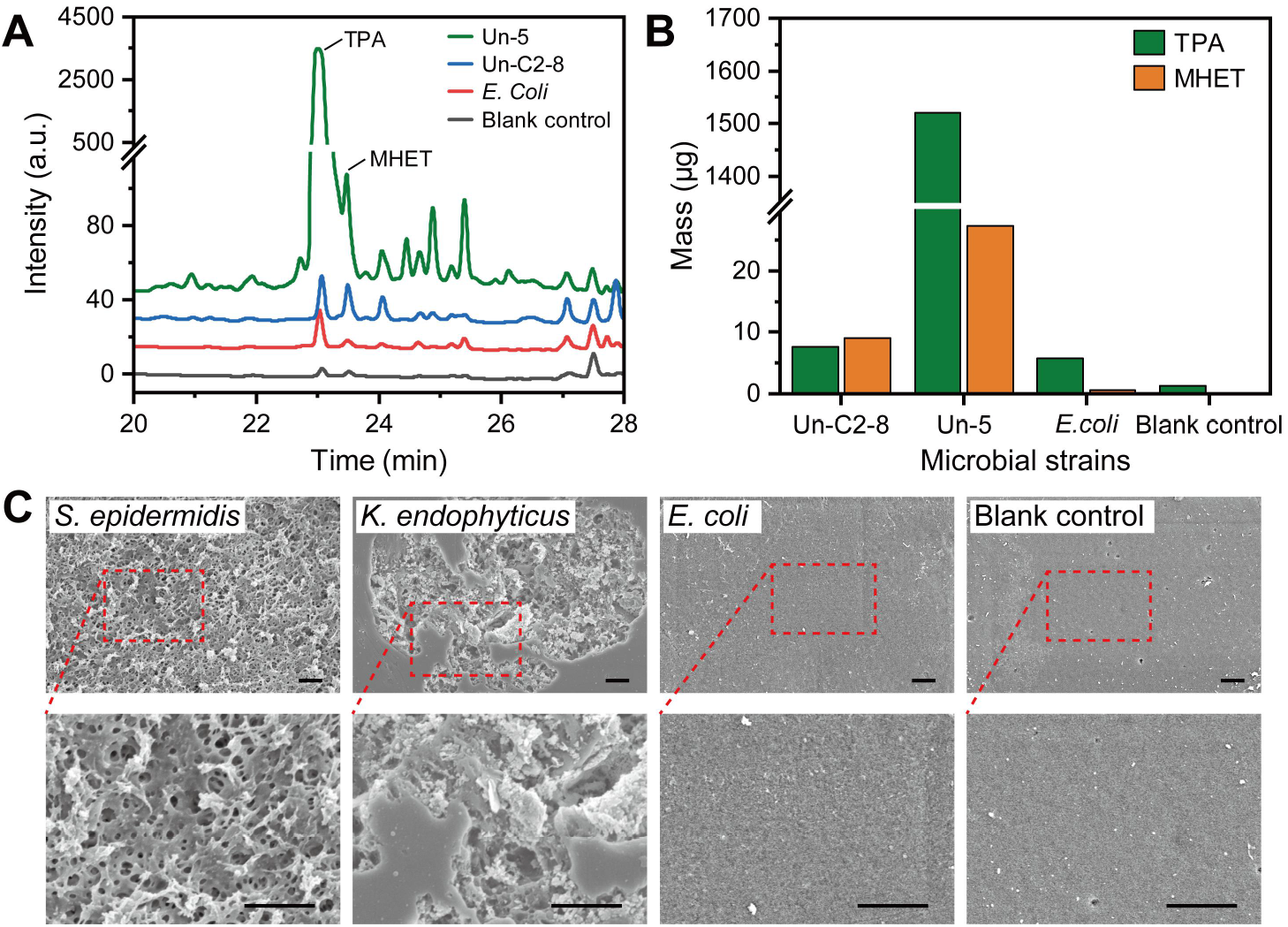
Validation of PET-biodegradation by FADS-obtained isolates K. endophyticus Un-5 and S. epidermidis Un-C2-8. (A) HPLC spectra of the degrading products released from the PET film incubated with Un-5, Un-C2-8, E. coli, and the blank control. (B) Mass conversion of PET film to TPA and MHET detected in culture supernatants from different strains and the blank control. (C) SEM images of PET films after 40-days degradation experiments with different strains versus the blank control. (Scale bar: 1 μm)

### Characterization of Putative PETases

Two putative PETases were selected following sequence interpretation of whole-genome sequences of Un-5 and Un-C2-8 and database searching: S9_948 belonging to the carboxylesterase family found in the genomes of both strains and PHB belonging to the dienelactone hydrolase family in the genome of Un-5. For PHB, its amino acid sequence is compared with those of a carboxylesterase from *Cereibacter sphaeroides* (PDB:4FHZ)^26^, a carboxyl esterase from *Rhodobacter sphaeroides* 2.4.1 (PDB: 4FTW), a metagenome-derived esterase (PDB: 3WYD)^27^, and a metagenome-derived esterase LC-Est5 (**Fig. S8**). PHB has a putative 28-residue signal peptide at their N-termini suggesting that it is secretory proteins like LC-Est1 (AIT56387.1, 25-residue signal peptide). The three amino acid residues that form a catalytic triad of esterolytic/lipolytic enzymes are fully conserved as Ser163, Asp220 in PHB (Ser165, Asp216, His248 in 4FHZ; Ser117, Asp165), and His197 in 3WYD. A pentapeptide GxSxG motif containing a catalytic serine residue is also conserved as GFSNG (residues 161–165) in PHB (**Fig. S8**). For S9_948, as expected for an α/β-hydrolase, its sequence contains a conserved Gly-X-Ser-X-Gly motif (GQSAG), which includes the catalytic serine residue (**Fig. S9**). The esterase Cbotu_EstA from the anaerobe *Clostridium botulinum* ATCC 3502 (PDB: 5AH1) was found to hydrolyze the polyester poly(butylene adipate-co-butylene terephthalate) (PBAT)^28^.

We then carried out degradation assays using p-NPB, FDBz, and PET films as the substrates to evaluate the degrading activity of S9_948 and PHB. We tried to extract and purify the enzymes heterologously expressed in *BL21(DE3)* but failed due to their low expression levels. PHB might impose inhibitory effect on the growth of *BL21(DE3)*. Therefore, crude enzymes were used for the following enzymatic assays. As expected, crude S9_948 and PHB exhibited significant kinetic hydrolytic activity against FDBz versus the blank control (**Fig. 5A**). Crude PHB and S9_948 also showed strong hydrolytic activities of 134.34±32.91 U·L^-1^ and 265.79±3.72 U·L^-1^ against *p*-NPB, respectively (**Fig. 5B**). Moreover, the SEM images reveal that both S9-948 and PHB can induce surface erosion to PET films after incubation for two weeks (**Fig. 5C**). However, the HPLC results of the relevant supernatants showed that hydrolysis products including BHET, MHET, and TPA, are below the detection limits. This result suggests that the enzymes might have low thermostability during long-term incubation. In addition, the biodegradation of PET by Un-5 and Un-C2-8 might involves multiple enzymes and steps and S9_948 and PHB are potentially participating enzymes. Considering the complexity of the PET biodegradation process, we expect that other key PETases are yet to be discovered for the microbial isolates obtained by FADS.

**Fig. 5.**
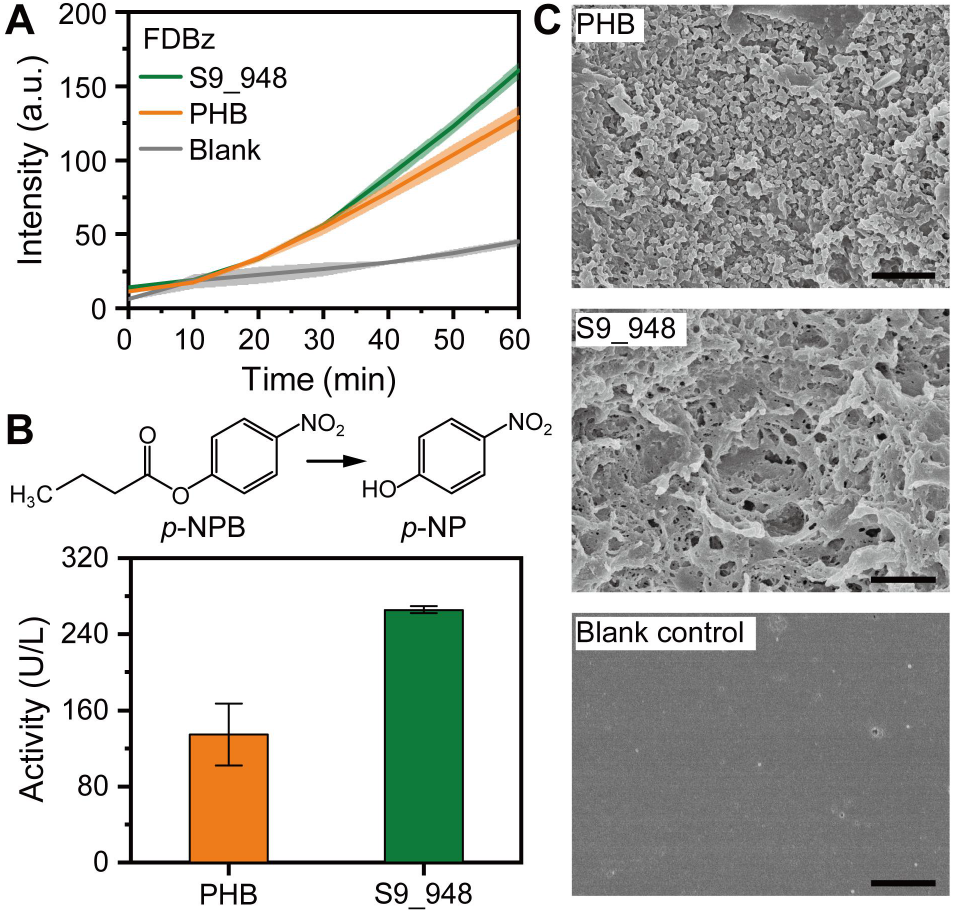
Enzymatic activity analysis of PHB and S9_948. (A) Hydrolysis of p-NPB measured by absorbance. (B) Hydrolysis of FDBz measured by fluorescence plate reader. (C) SEM images show surface erosion of PET films. (Scale bar: 1 μm)

## Discussion

In summary, we developed a complete FADS pipeline using FDBz as the fluorogenic probe for screening of single microbial cells at a large scale to advance the discovery of PET-degrading enzymes for biodegradation and sustainable recycling of PET. Because environmental microbes require longer incubation, we developed and optimized the pipeline to include long-term pre-incubation following microfluidic picoinjection of the fluorescence substrate FDBz. We then applied the system to screen an environmental sample from a PET textile mill and successfully obtained putative PET-degrading microbes belonging to nine different genera. Moreover, two putative PETases from environmental microbial strains K. endophyticus Un-5 and S. epidermidis Un-C2-8 were genetically derived, heterologously expressed, and preliminarily validated for their PET-degrading activity. Overall, the FADS pipeline opens possibilities for obtaining novel microorganisms and enzymes for PET biodegradation with superior throughput, sensitivity, and specificity. The FADS pipeline allows the directed evolution of PETases by random mutagenesis and screening. It also enables the screening of environmental microbes to recover novel enzymes that are not previously known or studied.

Our results prove that the FADS method could be applied to and effectively speed up the screening of environmental microbes present in various environments. This methodology may be extended to screen enzymes that degrade other synthetic plastics such as polyesters^29-30^, polyethylene^31-32^, and polycarbonate^33^ with the further development of specific fluorogenic probes targeting various synthetic plastics. Nano/micro-scale polymer particles coupled with fluorogenic enzymatic activity sensors are preferred to mimic the crystalline structures of plastic polymers in future work. To implement the FADS system in standard microbiology laboratories, we will focus on streamlining the instrument setup, device fabrication, and process automation.

## Materials and Methods

### Preparation of Microbial Samples

Wastewater samples were collected from a PET textile mill located in Southeast China (Shaoxing, Zhejiang, China) and stored at 4°C until use. We diluted each 10 g sample in 90 mL 1X PBS (pH 7.0) in a 250 mL flask and shook it at room temperature at 200 RPM for 30 min. Next, 1 mL diluted sample solution was added into PET-YSV medium (PET fiber 6 g/L, (NH_4_)_2_SO_4_ 0.2%, Trace elements 10%) for either direct FADS or FADS after enrichment cultivation. For enrichment cultivation, trace elements were added containing 0.1% FeSO_4_·7H_2_O, 0.1% MgSO_4_·7H_2_O, 0.01% CuSO_4_·H_2_O, 0.01% MnSO_4_·H_2_O and 0.01% ZnSO_4_·7H_2_O. The samples were cultivated for seven days at 37°C before FADS. Live-cell numbers of the original sample or its enrichment were adjusted to similar levels via live/dead staining and cell counting (LIVE/DEAD^®^ *Bac*Light™ Bacterial Viability kit, Molecular Probes, Eugene, OR, USA), and diluted to ∼7×10^7^ CFU·mL^-1^ in YSV medium to yield an average number of cells per droplet (*λ*) of 0.28 for single-cell sorting experiments.

### Microfluidic Devices Fabrication and Operations

The optical setup for the FADS experiment was built based on an inverted microscope (IX81, Olympus, Japan)^20^. Here, a 20 mW, 473 nm solid-state laser was shaped through a 20X objective into a 20 µm size spot focusing in the sorting channel. The fluorescence of the droplets was captured by the objective and split between a high-speed camera and a photomultiplier tube (10722-210, Hamamatsu Photonics). The signal output from the PMT was received and processed using a program written in LabVIEW (National Instrument, Austin, Texas, USA). Droplet sorting was triggered by a train of 1000-V, 30 kHz pulses applied by a high voltage amplifier (Trek). Polydimethylsiloxane (PDMS) was purchased from Momentive Performance Materials (Waterford, NY). Microfluidic devices were designed and fabricated as previously described^20^. Gastight glass syringes (Agilent, Reno, NV) were used for loading and infusing solutions into the devices by Syringe Pumps (Pump 11 PicoPlus, Harvard Apparatus, USA). Three devices, including the droplet maker, the picoinjector, and the droplet sorter were operated step-by-step as follows: **(1)** Droplet maker: microbial suspension samples were diluted to ∼7×10^7^ CFU/mL in YSV medium and infused into droplet maker device to generate single-cell encapsulating droplets (∼4 pL) at a throughput of 2700 drops per second. The droplets were collected in a 20-cm long tubing and incubated for three days to produce PET-degrading enzymes by positive microbial species. **(2)** Picoinjection: a solution of FDBz was introduced into the droplets by picoinjection and incubated for two hours. **(3)** Droplet sorter: the droplets were sorted based on the fluorescent intensity at a throughput of ∼1000 drops per second for several hours (∼3,600,000 droplets per hour), and those droplets with fluorescence above the sorting threshold were collected in an Eppendorf tube followed by demulsification and recovery of microbial cells for cultivation and enzymatic assays on agar plates. The fluorescence images of droplets were collected by an inverted fluorescence microscope (Eclipse Ti, Nikon, Tokyo, Japan).

### High-throughput Sequencing and Phylogenetic Analysis

Bacterial single colonies were picked from agar plates and inoculated in PET-YSV medium for subculture. DNA extractions from the isolates were carried out by using BMamp Rapid Bacterial DNA extraction kit (Cat. No. DL111-01, Biomed, Beijing, China) following the manufacturer’s instructions. Following PCR-amplified 16S rRNA gene sequencing, we obtained 16S rRNA identity of the strains using EzBioCloud (https://www.ezbiocloud.net/). Next, we aligned the sequences using ClustalW and plotted the phylogenetic tree with MEGA6 software using the neighbor-joining method^34^. For whole-genome sequencing of pure isolates, DNA extractions were sequenced on an Illumina HiSeq PE150 platform (Novogene Co., Ltd. Beijing, China).

### PET-degradation Activity Evaluation by clear zone formation and HPLC

The resulting microbial isolates were cultured in YSV medium added with PET fibers for three days, followed by clear zone formation in a solid media plate assay. The BHET-Basal agar plates were prepared with 0.4% BHET and 25mM Tris-HCl (pH 7.4) in 1.5% agar. The BHET-LB agar plates were prepared with 0.4% BHET and LB broth in 1.5% agar. Aliquots of 100 μL bacterial suspension from the YSV culture were added in punched holes on these agar plates and incubated at 37°C; the sizes of clear zones around the punched holes were measured after three days. Reverse-phase high-performance liquid chromatography (HPLC) was performed on an Agilent 1200 system (Agilent, Reno, NV) with a UV detector and a C18 column (Inertsil ODS-3, Shimadzu, Japan) as previously described^3^. The operation was performed using a gradient (methanol-20 mM phosphate buffer) that increased from 25–100% at 15–25 min with a flow of 1 mL·min^-1^. The injection volume was 10 μL, and the detection wavelength was 240 nm.

### Physicochemical and Morphological Characterization of PET Materials

A simultaneous thermal analyzer (NETZSCH STA 449F3, Germany) was used to perform thermogravimetric analysis (TGA) and differential thermal analysis (DTA) of PET fibers and films by heating 10–30 mg samples in an aluminum pan from 50 to 550°C at 20 K·min^-1^. The crystal structures of PET samples were quantified using an X-ray powder diffractometer (XRD) (SmartLab, Rigaku, Japan). For SEM imaging, the PET films were cleaned, air-dried, and coated by gold sputtering and observed under a SU8010 scanning electron microscope (SEM, Hitachi, Japan) at 5 kV to reveal surface degradation structures.

### Heterologous expression and preliminary activity validation of enzymes

The genes encoding putative PET-degrading enzymes, including S9_948 and PHB from *Kineococcus endophyticus* Un-5 and *Staphylococcus epidermidis* Un-C2-8 were selected for molecular cloning and heterologous expression. A set of specific primers were designed for PCR amplification of the target genes (Table 1) using the following steps. An initialization step of 5 min at 94°C was used to activate the Taq polymerease. Next, a total 30 cycles of amplification were performed as follows: a DNA denaturation step of 10 s at 98°C, a primer annealing step of 10 s at 55°C, and a DNA extension step of 20 s at 72°C. Afterward, a final extension step was performed for 2 min at 72°C. The PCR products were purified by agarose gel electrophoresis and sequenced for confirmation. Afterward, the target gene was connected to the expression vector pET-28a (Novagen) and transformed to *Escherichia coli* (*E. Coli*) *BL21(DE3)*. The recombinant putative enzymes were expressed as N-terminal fusion to His-tag. The transformed *BL21(DE3)* was cultured at 37°C in 50-mL LB supplemented with 50 μg·mL^-1^ Kanamycin to reach an OD_600_ of 0.6. The enzyme expression was then induced at 16°C overnight with shaking at 160 RPM. Following the induction time, cells were harvested by centrifugation and resuspended in 5 mL standard buffer (50 mM Tris-HCl (pH 8.0), 50 mM NaCl). Resuspended cells were lysed by an ultrasonic homogenizer (scientz-IID, Scientz Biotech, Ningbo, China) for 20 min. The supernatant (soluble fraction) containing the crude enzymes was separated from cell debris by centrifugation at 14,000 × g for 10 min and stored at 4°C for further use.

The FDBz hydrolysis from crude S9_948 or PHB was performed by mixing 100 μL enzyme solution with 100 μL 250 μM FDBz. The kinetic assays were carried out using the EnSpire plate reader at 37°C within 60 min. The hydrolysis of *p-*NPB by crude S9_948 or PHB was measured as previously described with slight modifications^35^. Absorbance assays were carried out on the EnSpire plate reader, using 200 μL reactions composed of 20 μL crude enzymes and 10 μL 1 M *p-*NPB in Tris buffer (pH 8.0). The production of *p*-nitrophenol (*p*-NP) was monitored at 405 nm within 20 min at 37°C. One unit of activity (1 U) was defined as the enzymatic production of 1 μmol *p*-NP per min at 37°C. The degradation of PET film by the crude S9_948 or PHB was evaluated by incubating PET film with the enzyme for seven days at 37°C, and evaluated by morphological characterization as described above.

## End Matter

### Notes

The authors declare no conflict of interest.

This article contains supporting information online.

Materials, Synthesis of Fluorescein dibenzoate, XRD analysis of PET materials, Determination of PET decomposition, a list of putative PET-degrading microbial isolates, PCR primers for molecular cloning of putative PETases, and degradation analyses of putative microbial isolates and enzymes (PDF)

## Acknowledgments

We thank Prof. Bian Wu from the Institute of Microbiology Chinese Academy of Sciences and Dr. Wenping Wu from Novozymes (China) Investment Co., Ltd. for their helpful discussions. This study was supported by the National Natural Science Foundation of China (21822408, 91951103, 31970091, 52073004), the Key Program of Frontier Sciences of the Chinese Academy of Sciences (QYZDB-SSW-SMC008), the National Key Research and Development Program of China (2018YFC0310703), Beijing Key Laboratory of Quality Evaluation Technology for Hygiene and Safety of Plastics (PQETGP2020003), and Novozymes A/S (Denmark).

## References

(1) Ammendolia, J.; Saturno, J.; Brooks, A. L.; Jacobs, S.; Jambeck, J. R.; An emerging source of plastic pollution: Environmental presence of plastic personal protective equipment (PPE) debris related to COVID-19 in a metropolitan city. Environ. Pollut. 2021, 269, 116160.

(2) Geyer, R.; Jambeck, J. R.; Law, K. L.; Production, use, and fate of all plastics ever made. Sci. Adv. 2017, 3, e1700782.

(3) Yoshida, S.; Hiraga, K.; Takehana, T.; Taniguchi, I.; Yamaji, H.; Maeda, Y.; Toyohara, K.; Miyamoto, K.; Kimura, Y.; Oda, K.; A bacterium that degrades and assimilates poly(ethylene terephthalate). Science 2016, 351, 1196–1199.

(4) Ribitsch, D.; Acero, E. H.; Greimel, K.; Eiteljoerg, I.; Trotscha, E.; Freddi, G.; Schwab, H.; Guebitz, G. M.; Characterization of a new cutinase from Thermobifida alba for PET-surface hydrolysis. Biocatalysis 2012, 30, 2–9.

(5) Hu, X.; Thumarat, U.; Zhang, X.; Tang, M.; Kawai, F.; Diversity of polyester-degrading bacteria in compost and molecular analysis of a thermoactive esterase from Thermobifida alba AHK119. Appl. Microbiol. Biotechnol. 2010, 87, 771–779.

(6) Carniel, A.; Valoni, R.; Junior, J. N.; Gomes, A. D. C.; Castro, A.; Lipase from Candida antarctica (CALB) and cutinase from Humicola insolens act synergistically for PET hydrolysis to terephthalic acid. Process Biochem. 2017, 59, 84–90.

(7) Costa, A.; Lopes, V.; Vidal, L.; Nicaud, J. M.; Coelho, M.; Poly(ethylene terephthalate) (PET) degradation by Yarrowia lipolytica: Investigations on cell growth, enzyme production and monomers consumption. Process Biochem. 2020, 95, 81–90.

(8) Koshti, R.; Mehta, L.; Samarth, N.; Biological Recycling of Polyethylene Terephthalate: A Mini-Review. J. Polym. Environ. 2018, 26, 3520–3529.

(9) Cui, Y.; Chen, Y.; Liu, X.; Dong, S.; Wu, B.; Computational Redesign of a PETase for Plastic Biodegradation under Ambient Condition by the GRAPE Strategy. ACS Catal. 2021, 11, 1340–1350.

(10) Sagong, H. Y.; Seo, H.; Kim, T.; Son, H. F.; Kim, K. J.; Decomposition of PET film by MHETase using Exo-PETase function. ACS Catal. 2020, 10, 4805–4812.

(11) Tournier, V.; Topham, C. M.; Gilles, A.; David, B.; Folgoas, C.; Moya-Leclair, E.; Kamionka, E.; Desrousseaux, M. L.; Texier, H.; Gavalda, S.; Cot, M.; Guemard, E.; Dalibey, M.; Nomme, J.; Cioci, G.; Barbe, S.; Chateau, M.; Andre, I.; Duquesne, S.; Marty, A.; An engineered PET depolymerase to break down and recycle plastic bottles. Nature 2020, 580, 216–219.

(12) Shinozaki, Y.; Watanabe, T.; Nakajima-Kambe, T.; Kitamoto, H. K.; Rapid and simple colorimetric assay for detecting the enzymatic degradation of biodegradable plastic films. J. Biosci. Bioeng. 2013, 115, 111–114.

(13) Thumarat, U.; Nakamura, R.; Kawabata, T.; Suzuki, H.; Kawai, F.; Biochemical and genetic analysis of a cutinase-type polyesterase from a thermophilic Thermobifida alba AHK119. Appl. Microbiol. Biot. 2012, 95, 419–430.

(14) Wei, R.; Oeser, T.; Schmidt, J.; Meier, R.; Barth, M.; Then, J.; Zimmermann, W.; Engineered bacterial polyester hydrolases efficiently degrade polyethylene terephthalate due to relieved product inhibition. Biotechnol. Bioeng. 2016, 113, 1658–1665.

(15) Zhong-Johnson, E.; Voigt, C. A.; Sinskey, A. J.; An absorbance method for analysis of enzymatic degradation kinetics of poly(ethylene terephthalate) films. Sci. Rep. 2021, 11, 928.

(16) Markel, U.; Essani, K. D.; Besirlioglu, V.; Schiffels, J.; Streit, W. R.; Schwaneberg, U.; Advances in ultrahigh-throughput screening for directed enzyme evolution. Chem. Soc. Rev. 2020, 49, 233–262.

(17) de Rond, T.; Danielewicz, M.; Northen, T.; High throughput screening of enzyme activity with mass spectrometry imaging. Curr. Opin. Biotech. 2015, 31, 1–9.

(18) Obexer, R.; Pott, M.; Zeymer, C.; Griffiths, A. D.; Hilvert, D.; Efficient laboratory evolution of computationally designed enzymes with low starting activities using fluorescence-activated droplet sorting. Protein Eng. Des. Sel. 2016, 29, 355–366.

(19) Baret, J. C.; Miller, O. J.; Taly, V.; Ryckelynck, M.; El-Harrak, A.; Frenz, L.; Rick, C.; Samuels, M. L.; Hutchison, J. B.; Agresti, J. J.; Link, D. R.; Weitz, D.; Griffiths, A. D.; Fluorescence-activated droplet sorting (FADS): efficient microfluidic cell sorting based on enzymatic activity. Lab Chip 2009, 9, 1850–1858.

(20) Qiao, Y.; Zhao, X.; Zhu, J.; Tu, R.; Dong, L.; Wang, L.; Dong, Z.; Wang, Q.; Du W; Fluorescence-activated droplet sorting of lipolytic microorganisms using a compact optical system. Lab Chip 2018, 18, 190–196.

(21) Vallejo, D.; Nikoomanzar, A.; Paegel, B. M.; Chaput, J. C.; Fluorescence-Activated Droplet Sorting for Single-Cell Directed Evolution. ACS Synth. Biol. 2019, 8, 1430–1440.

(22) Najah, M.; Calbrix, R.; Mahendra-Wijaya, I. P.; Beneyton, T.; Griffiths, A. D.; Drevelle, A.; Droplet-based microfluidics platform for ultra-high-throughput bioprospecting of cellulolytic microorganisms. Chem. Biol. 2014, 21, 1722–1732.

(23) Goto, H.; Kanai, Y.; Yotsui, A.; Shimokihara, S.; Shitara, S.; Oyobiki, R.; Fujiwara, K.; Watanabe, T.; Einaga, Y.; Matsumoto, Y.; Miki, N.; Doi, N.; Microfluidic screening system based on boron-doped diamond electrodes and dielectrophoretic sorting for directed evolution of NAD(P)-dependent oxidoreductases. Lab Chip 2020, 20, 852–861.

(24) Sjostrom, S. L.; Bai, Y.; Huang, M.; Liu, Z.; Nielsen, J.; Joensson, H. N.; Andersson, S. H.; High-throughput screening for industrial enzyme production hosts by droplet microfluidics. Lab Chip 2014, 14, 806–813.

(25) He, F. Y.; Wang, L. F.; Li, B. R.; Wang, Q. G.; Wang, Q.; Studies on crystal structure and hydrolysis feature of the fluorescein dibenzoate. Acta Chim. Sinica 1993, 51, 119–124.

(26) Ma, J.; Wu, L.; Guo, F.; Gu, J.; Tang, X.; Jiang, L.; Liu, J.; Zhou, J.; Yu, H.; Enhanced enantioselectivity of a carboxyl esterase from Rhodobacter sphaeroides by directed evolution. Appl. Microbiol. Biotechnol. 2013, 97, 4897–4906.

(27) Okano, H.; Hong, X.; Kanaya, E.; Angkawidjaja, C.; Kanaya, S.; Structural and biochemical characterization of a metagenome-derived esterase with a long N-terminal extension. Protein Sci. 2015, 24, 93–104.

(28) Perz, V.; Baumschlager, A.; Bleymaier, K.; Zitzenbacher, S.; Hromic, A.; Steinkellner, G.; Pairitsch, A.; Lyskowski, A.; Gruber, K.; Sinkel, C.; Kuper, U.; Ribitsch, D.; Guebitz, G. M.; Hydrolysis of synthetic polyesters by Clostridium botulinum esterases. Biotechnol. Bioeng. 2016, 113, 1024–1034.

(29) DelRe, C.; Jiang, Y.; Kang, P.; Kwon, J.; Hall, A.; Jayapurna, I.; Ruan, Z.; Ma, L.; Zolkin, K.; Li, T.; Scown, C. D.; Ritchie, R. O.; Russell, T. P.; Xu, T.; Near-complete depolymerization of polyesters with nano-dispersed enzymes. Nature 2021, 592, 558–563.

(30) Zadjelovic, V.; Chhun, A.; Quareshy, M.; Silvano, E.; Hernandez Fernaud, J. R.; Aguilo Ferretjans, M. M.; Bosch, R.; Dorador, C.; Gibson, M. I.; Christie Oleza, J. A.; Beyond oil degradation: enzymatic potential ofAlcanivorax to degrade natural and synthetic polyesters. Environ. Microbiol. 2020, 22, 1356–1369.

(31) Yang, J.; Yang, Y.; Wu, W. M.; Zhao, J.; Jiang, L.; Evidence of polyethylene biodegradation by bacterial strains from the guts of plastic-eating waxworms. Environ. Sci. Technol. 2014, 48, 13776–13784.

(32) Tennakoon, A.; Wu, X.; Paterson, A. L.; Patnaik, S.; Pei, Y.; LaPointe, A. M.; Ammal, S. C.; Hackler, R. A.; Heyden, A.; Slowing, I. I.; Coates, G. W.; Delferro, M.; Peters, B.; Huang, W.; Sadow, A. D.; Perras, F. A.; Ames Lab., I. U. S., Argonne National Lab. ANL, A. I. U. S., Catalytic upcycling of high-density polyethylene via a processive mechanism. Nat. Catal. 2020, 3.

(33) Artham, T.; Doble, M.; Biodegradation of physicochemically treated polycarbonate by fungi. Biomacromolecules 2010, 11, 20–28.

(34) Tamura, K.; Stecher, G.; Peterson, D.; Filipski, A.; Kumar, S.; MEGA6: Molecular Evolutionary Genetics Analysis version 6.0. Mol. Biol. Evol. 2013, 30, 2725–2729.

(35) Ate Lier, Z.; Metin, K.; Production and Partial Characterization of a Novel Thermostable Esterase from a Thermophilic Bacillus sp. Enzyme Microb. Tech. 2006, 38, 628–635.

