## Supplementary Materials for "Fluorescence-Activated Droplet Sorting of Polyethylene Terephthalate-degrading Enzymes"

Terephthalic acid (TPA, Aladdin), 4-nitrophenyl butyrate (*p*-NPB, Aladdin), and dimethyl sulfoxide (DMSO, Xilong Scientific) were acquired commercially. PET fibers, PET films, BHET, and MHET were provided by Novozymes A/S (Denmark). Fluorescein dibenzoate (FDBz) was synthesized by Novozymes (see supporting information for details)[34]. A 10-mM stock solution of FDBz in DMSO was prepared and diluted to the working concentration using 10 mM Tris-HCl (pH 7.4) for all experiments unless otherwise indicated. 'StickAway' cutinase and 'Lipex' lipase, both from Novozymes, were individually diluted in 10 mM Tris-HCl (pH 7.4) at a final concentration of 5 μM and used to evaluate the performance of FDBz as a PET-degrading indicator. Quantitative measurements of FDBz and *p*-NPB hydrolysis by enzymes or microbial strains were performed using a fluorescence reader (EnSpire multimode plate Reader, Perkin Elmer, Waltham, USA). Countess™ cell counting chamber slides (Thermo Fisher Scientific, MA, USA) were used for droplet array imaging under the microscope. Low melting-temperature liquid solder (Indium, Clinton, NY, USA) was used for making electrodes on microfluidic devices.

### Synthesis of Fluorescein dibenzoate (FDBz)

Fluorescein dibenzoate (formula I) was synthesized, which could be degraded by PETase and produces fluorescein (λex = 488 nm and λem = 523 nm). Briefly, fluorescein (400 mg, 1.2 mmol, Sigma-Aldrich) was suspended in toluene (30 ml, Sigma-Aldrich) to which benzoyl chloride (423 μL, 3.6 mmol, Sigma-Aldrich), triethyl amine (503 μL, 3.6 mmol, Sigma-Aldrich), and DMAP (5 mg, Sigma-Aldrich) were added. The mixture was stirred 24 h under $N_2$ at room temperature. Thin-layer chromatography (TLC, EtOAc-heptane 3:1) showed full conversion of fluorescein to FDBz. The mixture was purified by flash chromatography, eluted with a 0–100% gradient of EtOAc in heptane. The target compound was isolated in 605 mg (93% yield) and characterized by Nuclear Magnetic Resonance (NMR, Bruker) spectrums to confirm the molecular structure (**Fig. S1**, **Fig. S2**).

**XRD analysis of PET materials**

PET film and PET fiber were all measured to XRD, TGA, and DTA for thermal analysis (**Fig. S5**). XRD spectra showed the intensity of PET film and PET fiber in 2θ angle range from 10° to 70°, both exhibit three peaks at 2θ of about 18°, 23°, and 26°. It also indicated that PET fiber and film prepared have a similar crystal structure (**Fig. S5B**). TGA curves should be noted that relative mass losses occurred around 370-530°C. Alternatively, the areas of derivative thermogravimetric (DTG) were 81.8% and 78.6% for PET film and PET fiber, respectively (**Fig. S5C**). The thermal analysis results were displayed as DTA curves with temperature restricted to 50°C to 550°C (**Fig. S5D**). The curves were all characterized the similar thermal stability in both polymers. The large endothermic peaks appeared at 200-300°C, centered at 252°C for PET film and 265°C for PET fiber. The peak positions in DTA curves agree well for PET film and PET fiber, and it predicted the similarity in two polymer materials (**Fig. S5D**).

**Determination of PET decomposition products**

*Kineococcus endophyticus* Un-5 and *Staphylococcus epidermidis* Un-C2-8 were selected for PET-fiber biodegradation. HPLC curves clearly showed that *Kineococcus endophyticus* has a higher degrading performance than *Staphylococcus epidermidis* in the scale-up process. Determination of content of TPA and MHET and were 37.63±1.45 μg, 14.39±0.96 μg, 50.98±3.70 μg, and 16.20±7.54 μg by *K. endophyticus* and *S. epidermidis* in 20 mL PET-YSV medium, respectively (**Fig. S6**).

**Table S1**. Strains information and clear zone formation tests. The supernatants of liquid cultures of obtained strains were added into punched holes (8 mm in diameter) on the BHET-Basal or BHET-LB agar plates to perform clear zone formation assays.

| Strain ID | Top-hit taxon | Top-hit strain in EzBioCloud | Clear zone size on BHET-Basal (mm) | Clear zone size on BHET-LB (mm) |
|---|---|---|---|---|
| Un-C2-8 | *Staphylococcus epidermidis* | NCTC 11047 | 10.22 | 18.87 |
| Un-C2-9 | *Staphylococcus epidermidis* | NCTC 11047(T) | NT | NT |
| Un-C2-10 | *Staphylococcus epidermidis* | NCTC 11047(T) | NT | NT |
| Un-C2-11 | *Staphylococcus epidermidis* | NCTC 11047(T) | NT | NT |
| Un-4-5 | *Aeromonas caviae* | CECT 838 | NT | NT |
| Un-3-2 | *Pseudomonas protegens* | CHA0 | – | 11.4 |
| Un-5-5 | *Shewanella xiamenensis* | S4 | 14.07 | – |
| Un-5 | *Kineococcus endophyticus* | KLBMP 1274(T) | 13.69 | 19.93 |
| Un-10 | *Niveispirillum cyanobacteriorum* | TH16 | – | 10.1 |
| Un-14 | *Sinirhodobacter hankyongi* | BO-81 | – | 10.4 |
| PET-C2-1 | *Bacillus toyonensis* | BCT-7112(T) | NT | NT |
| PET-C2-7 | *Bacillus toyonensis* | BCT-7112 | 9.46 | – |
| PET-C2-2 | *Bacillus cereus* | ATCC 14579 | NT | NT |
| PET-C2-8 | *Bacillus cereus* | ATCC 14579 | 13.07 | – |
| PET-C2-W | *Bacillus cereus* | ATCC 14579 | NT | NT |
| PET-C2-6 | *Lysinibacillus fusiformis* | NBRC 15717 | NT | NT |
| PET-C2-A | *Lysinibacillus fusiformis* | NBRC 15717(T) | 8.8 | 10.75 |

Notes: '–', no clear zone formation; 'NT', not tested.

**Table S2.** The PCR primers for molecular cloning of putative PETases into the plasmid pET-28a(+). Primer sequences of target enzymes are underlined.

| Target | Primer | **Oligonucleotide sequence** |
|---|---|---|
| S9_948 | S9_948-F | TGCCGCGCGGCAGCCATATG<u>ATGGTTCAAGTCAAGATAGGTAACT</u> |
| | S9_948-R | TCGAGTGCGGCCGCAAGCTT<u>TTATTTATAGATATGTAACGCTTGA</u> |
| PHB | PHB-F | TGCCGCGCGGCAGCCATATG<u>GTGAGGGGTCGGGGGAGGGACGGTG</u> |
| | PHB-R | TCGAGTGCGGCCGCAAGCTT<u>TCAGGACGTCGCGGCCACCGACACG</u> |
| pET28a(+) | pET28a(+)-R | CATATGGCTGCCGCGCGGCA |
| | pET28a(+)-F | AAGCTTGCGGCCGCACTCGA |

### Supporting Figures

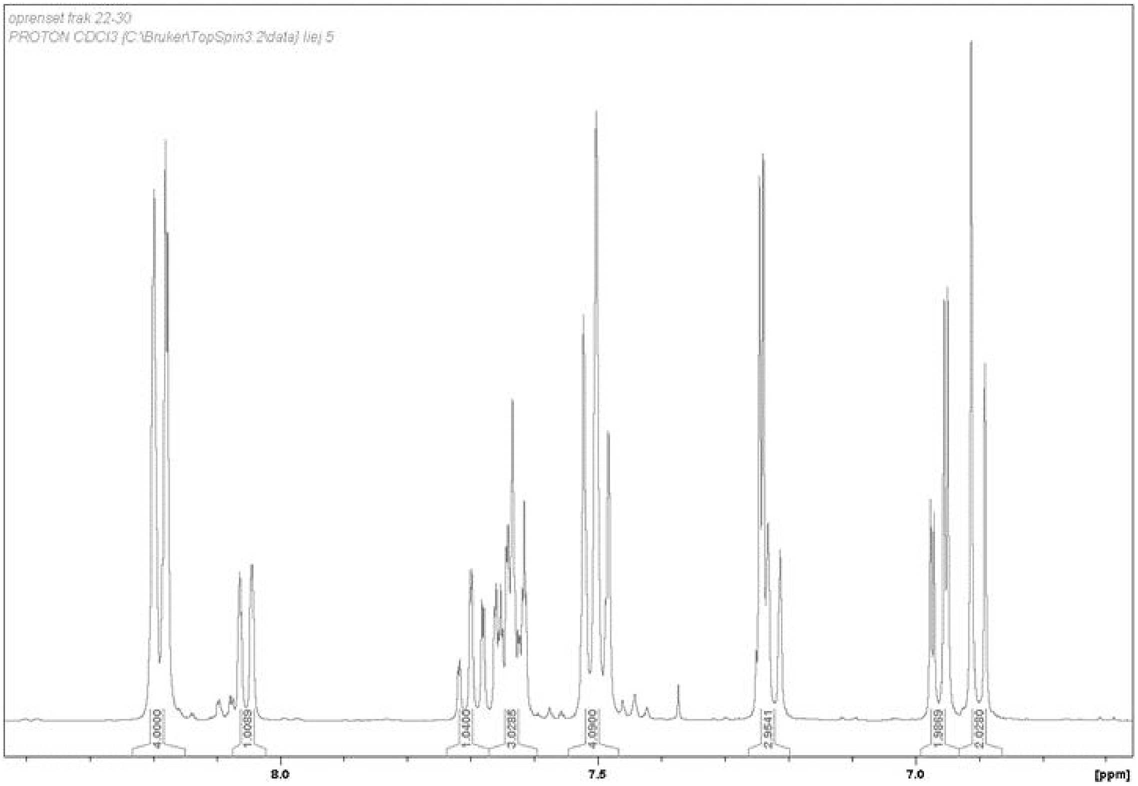


**Fig. S1.** $^1H$ NMR spectrum (400 MHz, $CDCl_3$) of FDBz: δ 8.24- 8.15 (m, 4H), 8.05 (d, 1H), 7.75-7.59 (m, 4H), 7.47-7.55 (t, 4H), 7.27-7.19 (m, 3H), 6.97-6.99 (dt, 4H).

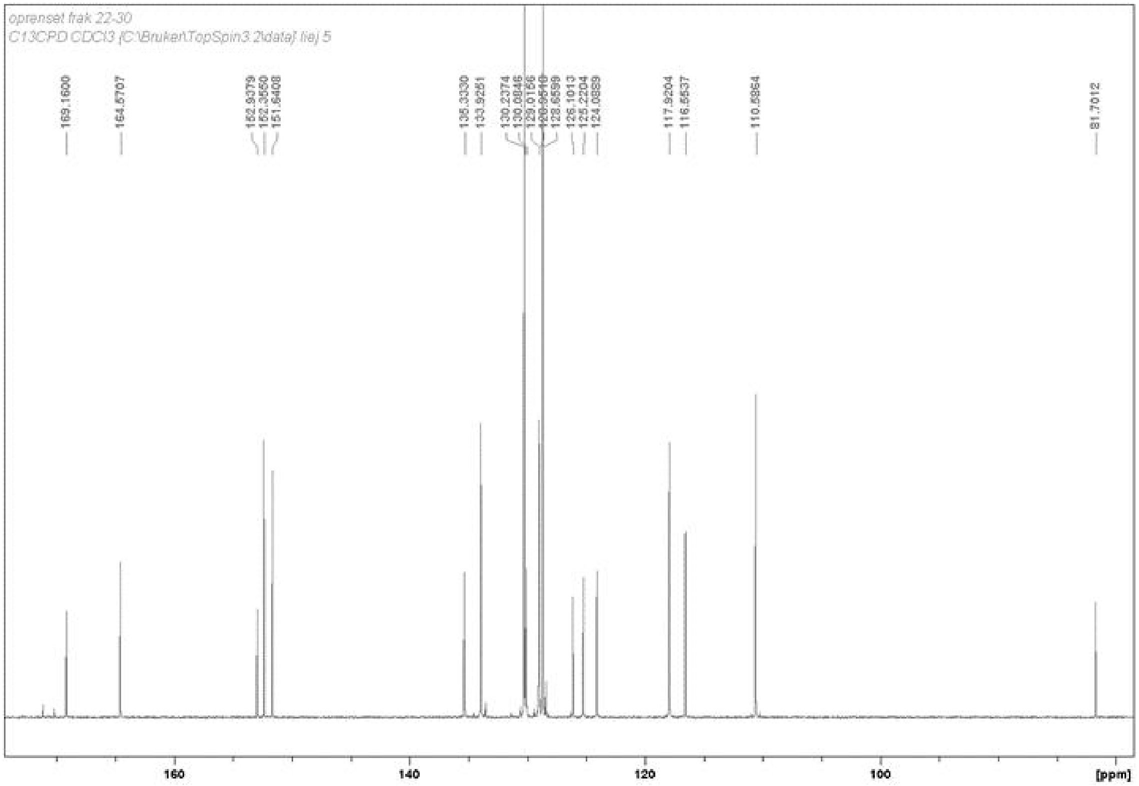


**Fig. S2.** $^{13}C$ NMR spectrum (100 MHz, $CDCl_3$) of FDBz: δ 169.21, 164.62, 152.99, 152.41, 151.70, 135.39, 134.59, 134.25, 133.98, 133.57, 130.63, 130.58, 130.29, 130.14, 130.03, 129.07, 129.01, 128.92, 128.71, 128.47, 126.32, 126.16, 125.28, 124.14, 117.98, 116.61, 110.64, 81.76.

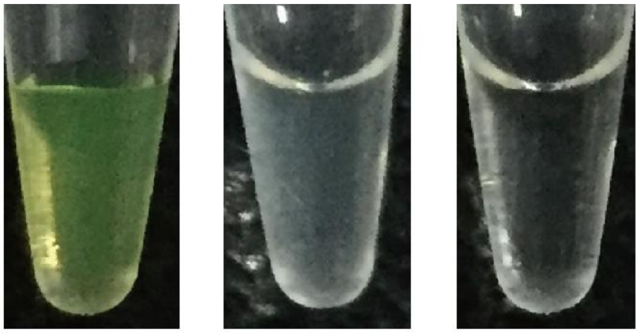


**Fig. S3.** The images show the chromogenic reaction of FDBz (250 μM) with cutinase (10 μM), lipase (10 μM), and blank control (Tris-buffer), all with a 1:1 mixing ratio.

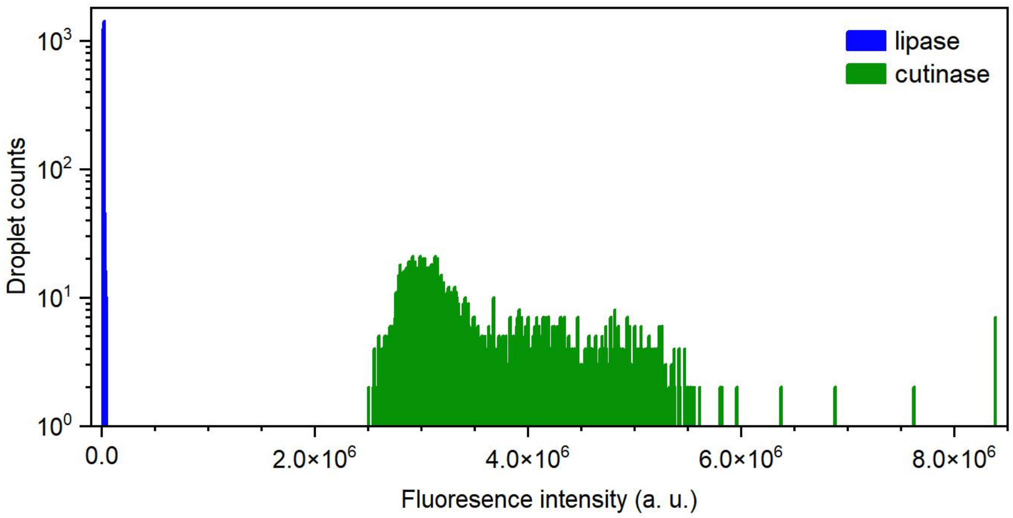


**Fig. S4.** The FADS data for 5 μM lipase or cutinase with 125-μM FDBz in 4-pL droplets after 2-h incubation (Green: cutinase; blue: lipase).

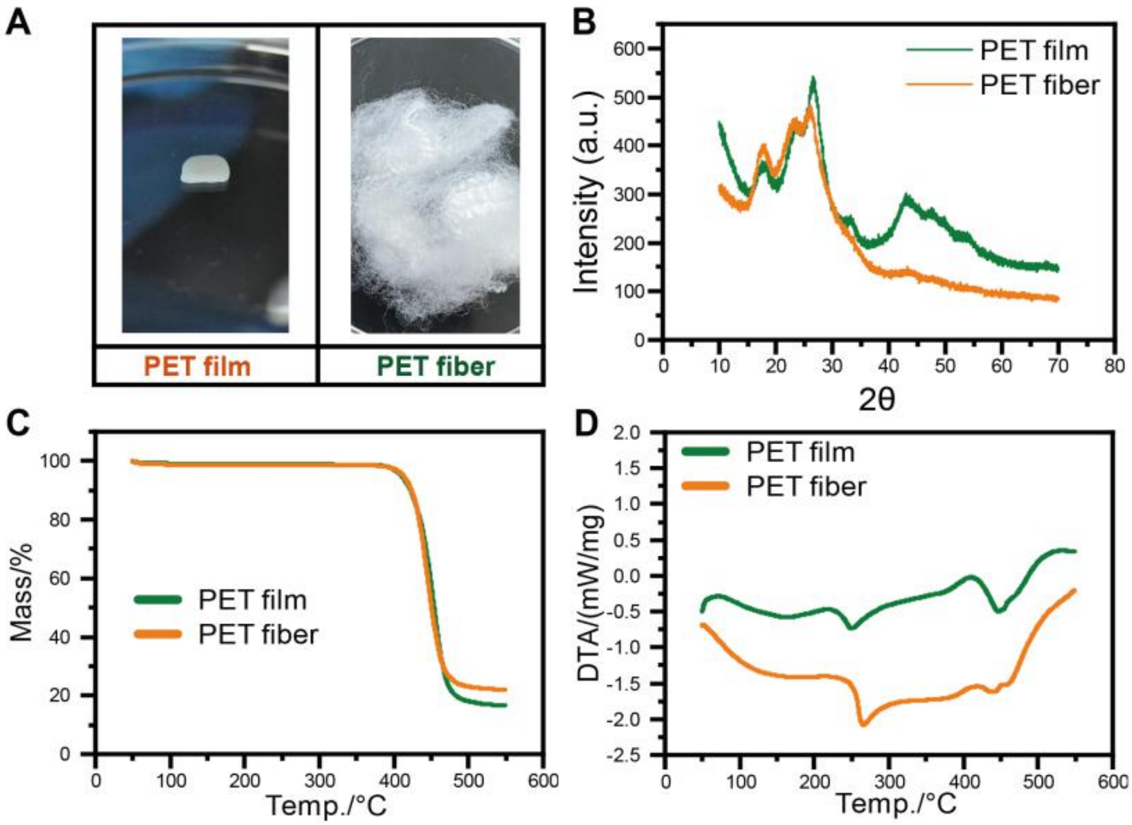


**Fig. S5.** Thermal analysis of PET films and PET fibers. (A) The pictures of PET films and pet fibers used in this study. (B) X-ray powder diffractometer (XRD) analysis. (C) thermogravimetric analysis (TGA). (D) Differential thermal analysis (DTA).

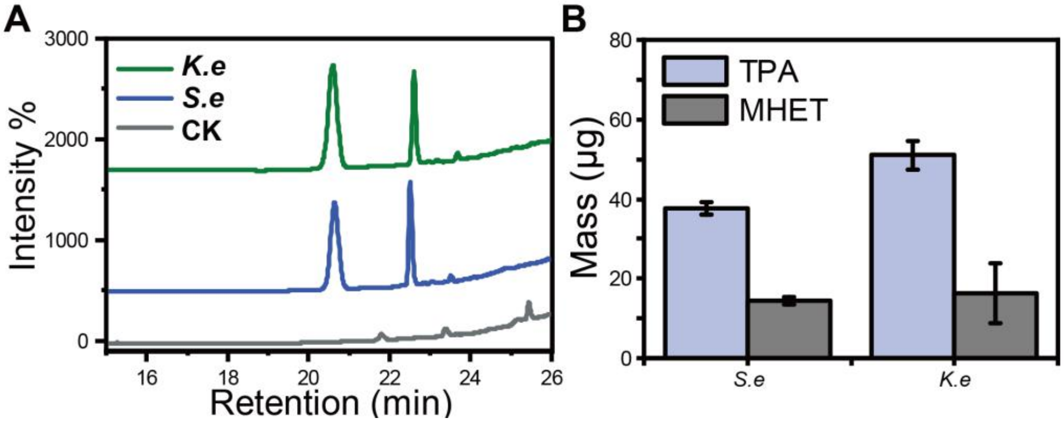


**Fig. S6.** (A) HPLC spectra of culture supernatants from strains Un-5 and Un-C2-8 supplemented with 60 mg PET fibers, accompanying with the blank control after two weeks. (B) Mass conversion of PET to TPA and MHET by the two strains in 20 mL supernatants after 14 days.

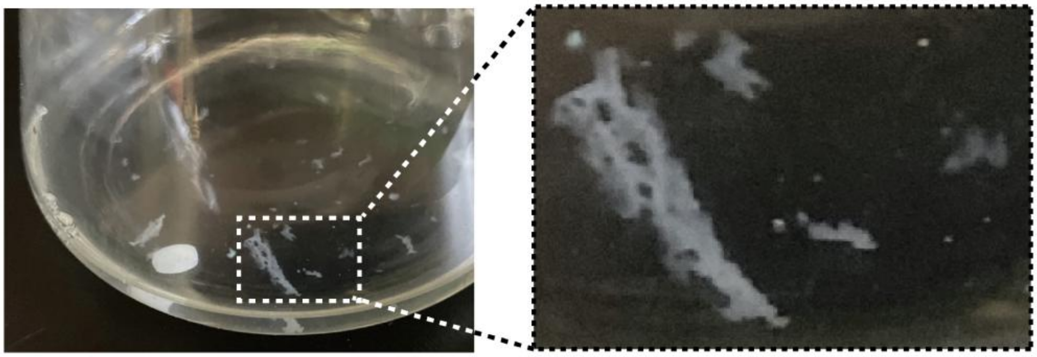

**Fig. S7.** Porous fragments break apart from PET films were observed after incubation with *Staphylococcus epidermidis* Un-C2-8 in 20 mL PET-YSV medium at 37°C for 40-days.

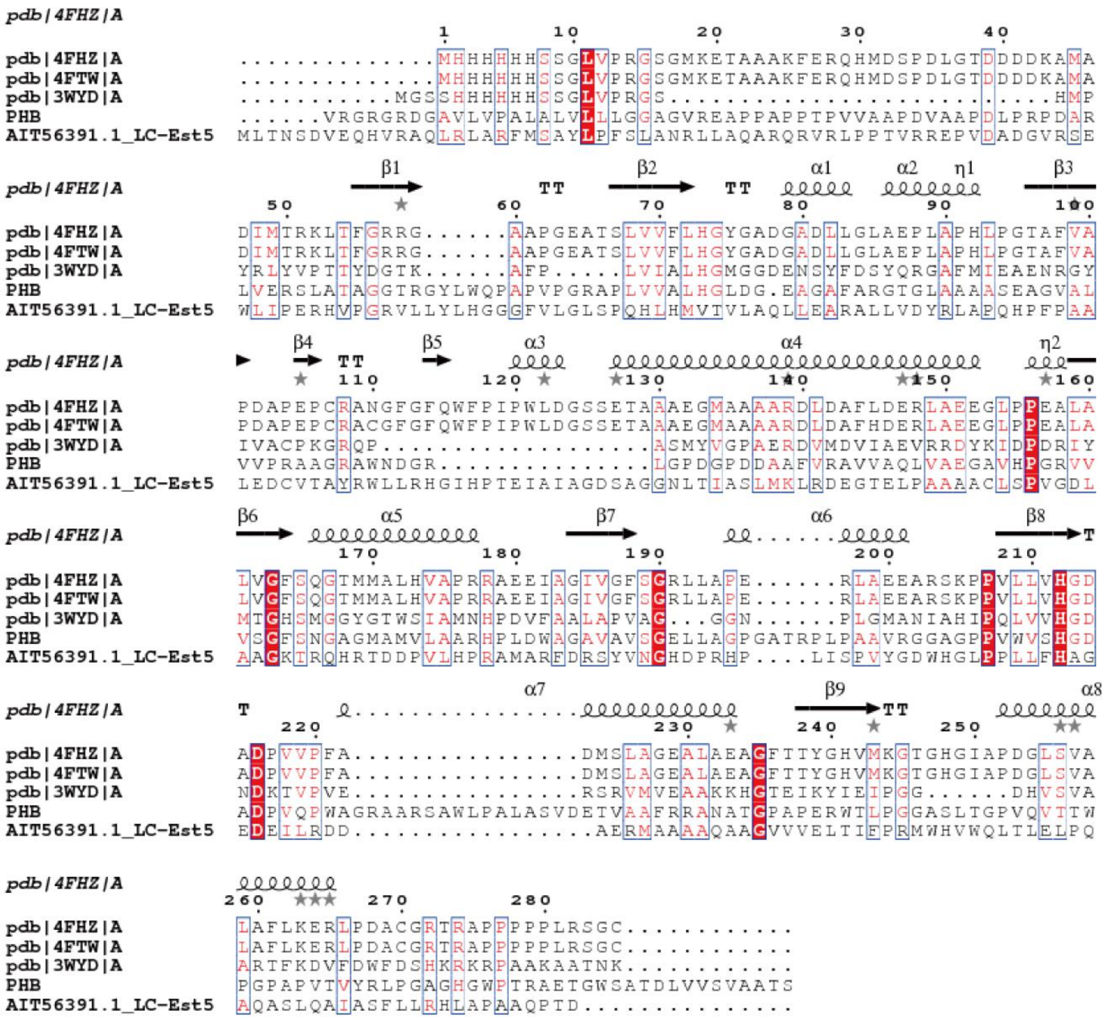


**Fig. S8.** Multiple sequence alignments of PHB with carboxyl esterase (PDB entry: 4FHZ and 4FTW), esterase (PDB entry: 3WYD), and Acetyl esterase/lipase (accession: AIT56391.1)

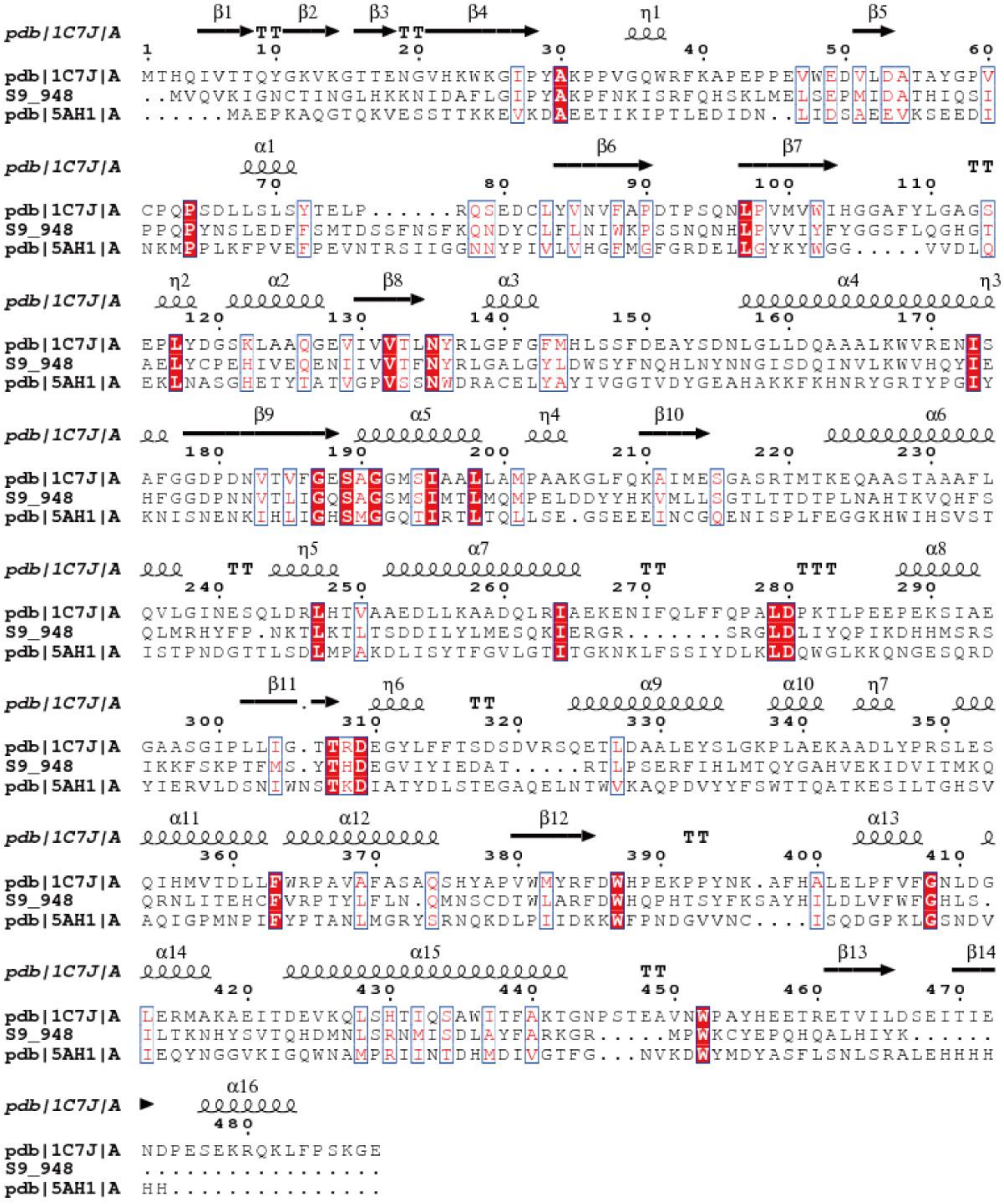


**Fig. S9.** Multiple sequence alignments of S9_948 with PNB esterase (PDB entry: 1C7J) and esterase (PDB entry: 5AH1).
